# Dissociable Roles of Ventromedial Prefrontal Cortex and Anterior Cingulate in Subjective Valuation of Prospective Effort

**DOI:** 10.1101/079467

**Authors:** Patrick S. Hogan, Joseph K. Galaro, Vikram S. Chib

## Abstract

The perceived effort level of an action shapes everyday decisions. Despite the importance of these perceptions for decision-making, the behavioral and neural representations of the subjective cost of effort are not well understood. While a number of studies have implicated anterior cingulate cortex (ACC) in decisions about effort/reward trade-offs, none have experimentally isolated effort valuation from reward and choice difficulty, a function that is commonly ascribed to this region. We used functional magnetic resonance imaging (fMRI) to monitor brain activity while human participants engaged in uncertain choices for prospective physical effort. Our task was designed to examine effort-based decision making in the absence of reward and separated from choice difficulty – allowing us to investigate the brain’s role in effort valuation, independent of these other factors. Participants exhibited subjectivity in their decision-making, displaying increased sensitivity to changes in subjective effort as objective effort levels increased. Analysis of blood-oxygenation level dependent (BOLD) activity revealed that the ventromedial prefrontal cortex (vmPFC) encoded the subjective valuation of prospective effort and ACC encoded choice difficulty. These results provide insight into the processes responsible for decision-making regarding effort, dissociating the roles of vmPFC and ACC in prospective valuation of effort and choice difficulty.

## INTRODUCTION

Our decisions are shaped not only by rewards, but also by the perceived level of physical effort required to obtain these rewards. Therefore the subjective perception of effort plays a critical role in driving choice – if the perceived effort of an action exceeds its subjective reward, the decision maker will optimally choose not to perform the action. Moreover, the perception of effort impacts everyday decisions in a variety of contexts ranging from job search (DellaVigna and Paserman 2005) and performing labor (Abeler et al. 2011; Augenblick et al. 2015), to participating in exercise and physical activity (Dishman 1991; Sniehotta et al. 2005). However, little is known about the neural representation of these subjective effort costs and how they are used by the brain to guide fundamental decisions to exert effort.

Subjective value signals for appetitive and aversive stimuli (e.g., money, food, aversive foods a liquids) have been found in the ventromedial prefrontal cortex (vmPFC) for both decision and outcome values (for comprehensive reviews of this literature see Bartra et al. 2013; Clithero and Rangel 2014; O’Doherty 2014). This body of work implicates vmPFC in the computation of subjective value across a multitude of stimuli with both positive and negative valence. Rather than experimentally separating such rewarding stimuli from effort costs, there have been a number of studies in animals (Walton et al. 2002; Rudebeck et al. 2008; Floresco et al. 2008; Walton et al. 2009; Hillman and Bilkey 2012) and humans (Croxson et al. 2009; Kurniawan et al. 2010; Prévost et al. 2010; Kurniawan et al. 2013; Skvortsova et al. 2014; Klein-Flügge et al. 2016; Chong et al. 2017) that examine how the brain makes decisions to trade effort for reward. This work suggests that anterior cingulate cortex (ACC) encodes the valuation of effort costs and effort cost trade-offs, both at the time of decision and at the time of effort exertion.

However, a recent series of studies, investigating neuroeconomic choice for prospective rewards, has shown that activity in ACC is best described by choice difficulty rather than valuation of the options presented (Shenhav et al. 2014; Shenhav et al. 2016). These experiments examined decisions regarding foraging costs and were careful to design their tasks such that prospective values were orthogonal to choice difficulty (i.e., the proximity in value between alternatives). From these studies it has been suggested that brain activity related to executive processing (e.g. choice difficulty) could confound valuation signals in ACC and throughout the brain in a variety of experimental contexts (Shenhav et al. 2014; Hayden and Heilbronner 2014; Westbrook and Braver 2015; Ebitz and Hayden 2016; Kolling et al. 2016b).

Thus, there are two potential limitations to the interpretation of previous work implicating a role for ACC in encoding effort costs: (i) effort costs were not experimentally isolated from reward; and (ii) effort costs may be correlated with choice. While a key feature of previous effort-based choice paradigms was that they examined decisions between effort and reward, they were not designed to experimentally isolate subjective effort costs from monetary or other reward-based incentives. As such, it is difficult to know if the identified neural signals are related to effort valuation per se, or multiplexed signals related to the context of effort-reward decision-making. Moreover, none of the previous studies of effort-based decision-making were designed to orthogonalize choice difficulty and effort value, which leads to the possibility that the ACC activity reported in these works could be related to the cognitive control associated with choice difficulty regarding decisions about effort and not effort valuation *per se*. In this study we attempted to address these limitations in the understanding of effort-based decision-making.

Here we investigated the behavioral representations of subjective effort cost and how these effort preferences are encoded in the brain’s valuation and decision-making circuitry. We had participants perform a novel effort-choice task while their neural activity was recorded with functional magnetic resonance imaging (fMRI). Our task did not involve monetary earnings, which allowed us to experimentally isolate effort cost (i.e., choices did not involve a trade-off between effort and reward). Furthermore, the effort choices presented were designed to experimentally separate choice difficulty and effort value to better understand the specific role ACC plays in effort-based decision-making. We hypothesized that, in a similar fashion to monetary rewards (Kahneman and Tversky 1979; Holt and Laury 2002), participants would exhibit subjectivity in their decisions for prospective effort. That is, effort representations would differ from the objective amount of effort, and would instead contain a degree of subjectivity driven by an individual’s perception of how effortful a task feels. Since our paradigm was designed to isolate subjective effort costs, independent of reward and choice difficulty, we hypothesized that behavioral representations of this effort subjectivity would be encoded in vmPFC – consistent with findings in human and animal studies investigating subjective value of appetitive and aversive stimuli. Additionally, we hypothesized that when choice difficulty and effort value were experimentally isolated, ACC would encode the former in concert with previous studies of neuroeconomic choice.

## MATERIALS AND METHODS

### Experimental Setup

Presentation of visual stimulus and acquisition of behavioral data were accomplished using custom MATLAB (http://www.mathworks.com) scripts implementing the PsychToolBox libraries (Brainard 1997). During functional magnetic resonance imaging (fMRI), visual feedback was presented via a projector positioned at the back of the room. Participants viewed a reflection of the projector in a mirror attached to the scanner head coil.

An MRI compatible hand clench dynamometer (TSD121B-MRI, BIOPAC Systems, Inc., Goleta, CA) was used to record grip force effort. During experiments, signals from this sensor were sent to our custom designed software for visual real-time feedback of participants’ effort exertion. Effort exertion was performed while participants held the force transducer in their right hand with arm extended while lying in the supine position.

To record participants’ choices we used an MRI compatible multiple button-press response box (Cedrus RB-830, Cedrus Corp., San Pedro, CA).

### Experiment Procedures

#### Participants

All participants were right handed, and were prescreened to exclude those with prior history of neurological or psychiatric illness. The Johns Hopkins School of Medicine Institutional Review Board approved this study, and all participants gave informed consent.

Thirty-eight healthy participants participated in the experiment, eight of which, were excluded for one or a combination of behavioral reasons. First, participants were excluded if they were unable to generate salient associations between effort levels and applied effort (*n* = 4; r-squared value between reported effort and perfect reporting less than 0.5). Second, inconsistent decision-making precluded participants from subsequent analyses (*n* = 5; random or near random choices, characterized by a temperature parameter less than 0.005). The final analyses included *n* = 30 participants in total (mean age, 23 years; age range, 18-34 years; 13 females).

#### Effort Paradigm

Prior to the experiment, participants were informed that they would receive a fixed show-up fee of $30. It was made clear that this fee did not, in any way, depend on performance or behavior over the course of the experiment.

The experiment began with acquiring participants’ maximum voluntary contraction (MVC) by selecting the maximum force achieved over the course of three consecutive repetitions on the hand clench dynamometer. During these repetitions participants did not have knowledge about the subsequent experimental phases, and were instructed and verbally encouraged to squeeze with their maximum force.

Next, participants performed an association phase in which they were trained to associate effort levels (defined relative to MVC) with the force they exerted against the hand dynamometer (Fig. 1A). Effort levels existed on a scale that ranged from 0 (corresponding to no exertion) to 100 (corresponding to a force equal to 80% of a participant’s MVC). A single training block consisted of five trials of training for each target level, where the target levels varied from 10 to 80 in increments of 10, and training blocks were presented in a randomized order. We did not perform association trials at the highest levels of effort to minimize the possibility that participants would become fatigued during this phase. A single trial of a training block began with the numeric display of the target effort level (2 s), followed by an effort task with visual-feedback in the form of a black vertical bar, similar in design to a thermometer, which increased in white the harder participants gripped the dynamometer (4 s). The bottom and top of this effort gauge represented effort levels 0 and 100, respectively. Participants were instructed to reach the target zone (defined as ±5 effort levels of the target) as fast as possible and maintain their force within the target zone for as long as possible over the course of 4 seconds. Visual indication of the target zone was colored green if the effort produced was within the target zone, and red otherwise. At the end of the effort exertion, if individuals were within the target zone for more than two thirds of the total time (2.67 s) during squeezing, the trial was counted a success. These success criteria were meant to ensure that participants were exerting effort for a similar duration across all effort conditions. To minimize participants’ fatigue, a fixation cross (2-5 s) separated the trails within a training block and 60 seconds of rest were provided between training blocks.

**Figure 1.**
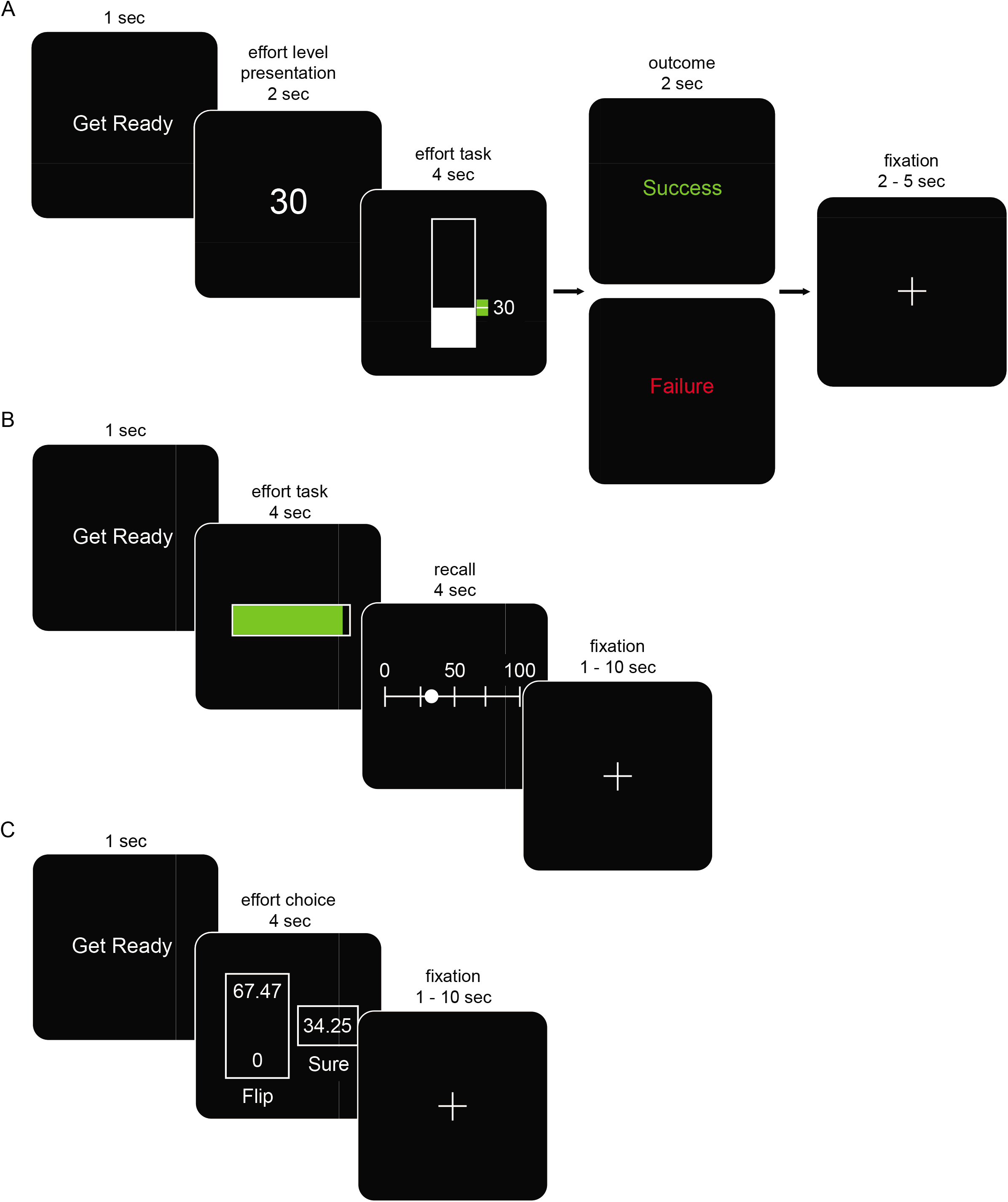
Experimental paradigm. A. Association phase; participants were trained to associate numeric effort levels with force exerted on a hand-clench dynamometer. Effort levels ranged from 0 (no force) to 100 (80% of maximum grip force). A training block consisted of five trials each at a series of target effort levels. Each trial began with presentation of the target, followed by an effortful grip with real-time visual feedback of the exerted force represented as a bar that increased in height with increased exertion. A green visual cue was also displayed, within which participants were instructed to maintain their exerted effort. Feedback of success or failure was provided at the end of each trial.
B. Recall phase; participants were instructed to fill a horizontal bar by gripping the transducer. On each trial, the full bar corresponded to a different target effort level that was unknown to participants. Successfully achieving the effort target resulted in the bar turning from red to green. Following exertion, participants used push-buttons to move a cursor along a 0−100 number line to select the effort level they believed they had squeezed. No feedback was provided as to the accuracy of participants’ reported effort levels.
C. Choice phase; participants were presented a series of risky gambles which involved choosing between two options: exerting a low amount of effort with certainty (“Sure”), or taking a gamble that could result in either a higher level of exertion or no exertion with equal probability (“Flip”). Gambles were not realized following a choice. At the end of the choice phase, to ensure participants revealed their true preferences for effort, 10 choices were randomly selected and played out such that any effort required would need to be exerted before they completed the experiment.

Following the association phase, we performed a recall phase to test if participants successfully developed an association between the effort levels and the actual effort exerted (Fig. 1B). Participants were tested on each of the previously trained effort levels (10 to 80, increments of 10), six times per level, presented in a random order. Each recall trial consisted of the display of a black horizontal bar that participants were instructed to completely fill by gripping the transducer – turning the force-feedback from red to green once the target effort level was reached. For the recall phase, the full bar did not correspond to effort level 100 as in the previous phase, but instead was representative of the target effort level being tested in a particular trial. Participants were instructed to reach the target zone as fast as possible, to maintain their produced force as long as possible, and to get a sense of what effort level they were gripping during exertion (4 s). Following this exertion, participants were presented a number line (from 0 to 100) and told to select the effort level they believed they had just gripped. Selection was accomplished by using two push-buttons to move a cursor left and right along the number line, and a third button to enter their believed effort level. Participants had a limited amount of time to make this effort assessment (4 s), and if no effort level was selected within the allotted time the trial was considered missed. No feedback was given to participants as to the accuracy of their selection.

Finally, during the choice phase of the experiment we scanned participants’ brains with fMRI while they were presented with a series of effort gambles and the choices from these gambles were used to characterize how individuals subjectively valued effort (Fig. 1C). Prior to being presented with the effort gambles participants were told that 10 of their decisions would be selected at random at the end of the experiment, and that they would have to remain in the testing area until they achieved the exertions required. This was done to ensure that participants were properly incentivized on each trial. Importantly, effort choices and realization occurred in separate sessions, which allowed us to examine the neural signals associated with prospective effort valuation unaffected by physical fatigue. A single effort gamble consisted of choosing between two options shown on the screen under a time constraint (4 s): one option entailed exerting a low amount of force (*S*) with certainty (known as the “sure” option); whereas the other entailed taking a risk which could result in either high exertion (*G*) or no exertion, with equal probability (known as the “flip” option). The effort levels were presented using the 0 to 100 scale that participants were trained on during the association phase. Participants made their choices by pressing one of two buttons on a hand-held button-box with their right hand with either their first or second digits. Gambles were not resolved following choice. Effort gambles (170 in total) were presented consecutively in a randomized order. Participants were encouraged to make a choice on every trial, however there was no penalty for failing to make a decision within the four second time window (average percent of missed trials across subjects 1.8% ± 3.8). Failure to make a choice on time was logged as a missed trial, and was not repeated.

At the end of the choice phase, the computer selected 10 of the trials at random to be implemented. The outcomes of the selected trials, and only those trials, were implemented. In this way, participants did not have to worry about spreading their effort exertion over all of their trials. Critically participants were instructed that the experiment would not be completed, and they were to remain in the testing area, until they achieved the exertions randomly implemented from the choice phase.

Our effort-based choice task has three properties that are important to stress. First, in our experimental design, we exploit the theoretical equivalence between risk preferences and subjective valuation in such a way that we can measure subjective valuation via the presentation of risky choices concerning effort, a widely accepted practice in economics and decision-neuroscience (Camerer et al. 2005; Rangel et al. 2008). Second, it is important to note that these effort choices did not involve monetary earnings or exertion at the time of choice, which allowed us to experimentally isolate prospective effort value representations from reward and fatigue. Third, effort choices were designed to span a range of potential effort values, capturing behavioral extremes of choice acceptance and rejection, centered at indifference. In doing so, this design ensures that choice difficulty, the magnitude of the relative value of effort options (behaviorally indexed by reaction time), is orthogonal to the difference in value between the options.

#### Effort Choice Values

The effort amounts were chosen to accommodate a range of effort sensitivity. For each trial, we denote the ratio η = G/S, of the worst possible outcome (choosing flip and having to exert the positive effort level) to the amount of effort in the sure option. We reasoned that participants would primarily exhibit increasing marginal utility for effort, and we therefore chose a range of η ∈ [1.75, 2.75]. In our gamble set, the force level associated with the sure option ranged from 5-35 in increments of 3.25 and were multiplied by the ratio η to generate 100 unique effort gambles, all with effort levels below 100. To span a broader range of η, additional gambles were designed in a similar method described above, except that the ratio of flip to sure (η) was halved and then multiplied by the sure values (η_1/2_ ∈ [0.88, 1.38]). 30 of these 100 additional effort gambles resulted in trivial (*G*,*S*) pairings with the flip values less than sure values (η < 1), and were thus excluded. The end result was 170 unique effort gambles (see Supplementary Materials for the full choice set).

#### Monetary Choice Task (Prospect Theory Task)

To investigate if there was a relationship between subjective preferences for effort and monetary gains and losses (i.e. risk aversion, loss aversion), a subset of participants (*n* = 22) performed a binary forced-choice task for money outside of the scanner. In this task participants made series of choices between a certain option involving a payout with 100% probability and a risky option involving gain and loss with equal probability. This exact paradigm has been used in a number of studies to elicit subjective preferences for monetary gains and losses (Sokol-Hessner et al. 2009; Frydman et al. 2011; Sokol-Hessner et al. 2012; Chib et al. 2014).

#### MRI Protocol

A 3 Tesla Philips Achieva Quasar X-series MRI scanner and radio frequency coil was used for all the MR scanning sessions. High resolution structural images were collected using a standard MPRAGE pulse sequence, providing full brain coverage at a resolution of 1 mm × 1 mm × 1 mm. Functional images were collected at an angle of 30° from the anterior commissure-posterior commissure (AC-PC) axis, which reduced signal dropout in the orbitofrontal cortex (Deichmann et al. 2003). Forty-eight slices were acquired at a resolution of 3 mm × 3 mm × 2 mm, providing whole brain coverage. An echo-planar imaging (FE EPI) pulse sequence was used (TR = 2800 ms, TE = 30 ms, FOV = 240, flip angle = 70°).

### Data Analysis

#### Effort Choice Analysis

We used a two parameter model to estimate participants’ subjective effort cost functions. We assumed a participant’s cost function *V*(*e*) for effort *e* as a power function of the form:

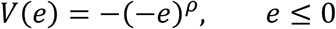

In this definition of effort cost, the effort level *e* is defined as negative, with the interpretation being that force production is perceived as a loss. The parameter *ρ* represents sensitivity to changes in subjective effort value as the effort level changes. A large *ρ* represents a high sensitivity to increases in absolute effort level. *ρ* = 1 implies that subjective effort costs coincide with objective effort costs.

Representing the effort levels as prospective costs, and assuming participants combine probabilities and utilities linearly, the difference in value between the two effort options relative to the sure prospect, can be written as follows:

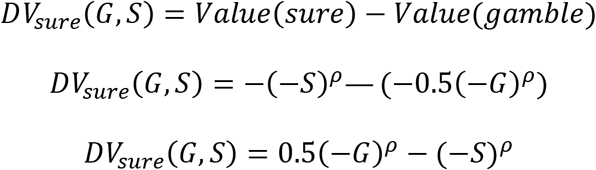

Where DV_sure_ denotes the difference in value between the two options, and both G < 0 and S < 0 for all trials.

We then assume that the probability that a participant chooses the sure option for the *k*^th^ trial is given by the softmax function:

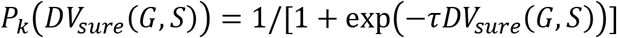

where *τ* is a temperature parameter representing the stochasticity of a participant’s choice (*τ* = 0 corresponds to random choice).

We used maximum likelihood to estimate parameters *ρ* and *τ* for each participant, using 170 trials of effort choices (*G*, *S*) with a participant’s choice denoted by *y* ∈ {0,1}. Here, *y* = 1 indicates that the participant chose the sure option. This estimation was performed by maximizing the likelihood function separately for each participant:

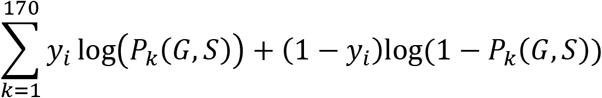

#### Monetary Choice Analysis

A separate maximum likelihood procedure was used to estimate parameters for monetary reward in a similar manner described in (Sokol-Hessner et al. 2009; Frydman et al. 2011; Chib et al. 2012; Sokol-Hessner et al. 2012), estimating both risk and loss aversion parameters for each participant. We expressed participants’ utility function *u* for monetary values *x* as

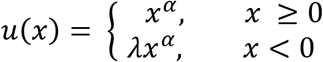

In this formulation, *λ* represents the relative weighting of losses to gains, and *α* represents the degree of a participant’s risk aversion. Assuming that participants combine probabilities and utilities linearly, the expected utility of a mixed gamble can be written as *U(G,L,S)* = (0.5 *G^α^* + 0.5 *λ*L) – *S^*α*^*, where G, L, S are the respective gain, loss, and sure options of the presented risky option.

The probability that a participant chooses the risky option for the *k*^th^ trial is given by the softmax function:

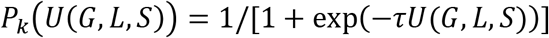

where *τ* is a temperature parameter representing the stochasticity of a participant’s choice.

The maximum likelihood procedure was accomplished using 140 gambles with participant response *y* ∈ {0,1}. Here, *y* = 1 indicates that the participant chose to make a gamble. The estimation was performed by maximizing the likelihood function:

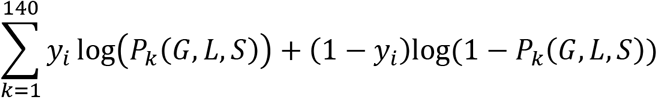

Mean and standard deviation for estimates are as follows: risk aversion: *α* = 0.81 (0.30), loss aversion: *λ* = 1.69 (1.32), temperature parameter: *τ* = 1.91 (1.14). Of the twenty-two included participants in the subjective effort experiment, four were excluded from this monetary analysis on the basis of inconsistent choices (*n* = 2) and parameter estimates beyond two standard deviations from the mean (*n* = 2). Exclusion from the analysis on these grounds was independent of the subjective effort experiment, and vice versa.

#### Image Processing and fMRI Statistical Analysis

The SPM12 software package was used to analyze the fMRI data (Wellcome Trust Centre for Neuroimaging, Institute of Neurology; London, UK). A slice-timing correction was applied to the functional images to adjust for the fact that different slices within each image were acquired at slightly different time-points. Images were corrected for participant motion, spatially transformed to match a standard echo-planar imaging template brain, and smoothed using a 3D Gaussian kernel (8 mm FWHM) to account for anatomical differences between participants.

To examine regions of the brain that encode participants’ subjective effort costs, we estimated participant-specific (first level) general linear models (GLM) for the effort choice phase of the experiment. This GLM included an event based condition at the time of effort choice, and parametric modulators corresponding to both a participant’s difference in subjective value between the gamble and sure options *DV_sure_*(*G*,*S*), and a behavioral measure for choice difficulty, log(response time). Trials with missing responses were modeled as a separate nuisance regressor. In addition, regressors modeling the head motion as derived from the affine part of the realignment procedure were included in the model. Using this model we were able to test brain areas in which activity was related to participants’ difference in subjective value between the gamble and sure effort options and their underlying subjective effort value representations, as well as activity related to choice difficulty. Using these conditions we created contrasts with the aforementioned parametric modulators, for difference in value, and choice difficulty at the time of effort choice.

We constructed a separate GLM incorporating a second, model-based measure of choice difficulty, -|DV_sure_|, as well as DV_sure_ and log(response time) as parametric modulators at the time of choice. With this additional model, we aimed to confirm that we still observed significant activation in ACC related to a model-based choice difficulty -|DV_sure_| after removing the variance associated with response time.

#### Statistical Inference

We analyzed the vmPFC signals shown within an independent region of interest (ROI) defined from an extensive meta-analysis of studies examining valuation of appetitive and aversive stimuli (5 mm radius sphere centered at Montreal Neurological Institute coordinates [2,46,−8]) (Bartra et al. 2013).

There is a degree of heterogeneity in the ACC activations reported in previous studies of effort-based decision making. With this in mind we analyzed the ACC signals shown within an independent ROI defined by averaging peak activations from a number of studies of effort-based decision making ([−6,−8,58] [4,−2,54] (Croxson et al. 2009); [6,24,28] (Prévost et al. 2010); [3,26,25] (Kurniawan et al. 2010); [6,23,28] (Kurniawan et al. 2013); [−6,4,42] (Skvortsova et al. 2014); [−6,11,34] (Klein-Flügge et al. 2016); [10,22,42] (Chong et al. 2017); mean coordinate values [1,12,39]).

For these ROIs, we regressed our design matrix on a representative time course, calculated as the first eigenvariate. This provides a very sensitive analysis because only a single regression is performed for this region and no multiple comparisons are required. The results of these ROI analyses were used for all statistical inferences about brain activity and are reported in the main text.

To clarify the signal pattern in each ROI we created plots of effect sizes for terciles (i.e., low, medium, high) at the peak of activity. It is important to note that these signals are not statistically independent (Kriegeskorte et al. 2009) and these plots were not used for statistical inference. They are shown solely for illustrative purposes.

#### Bayesian Model Selection of Imaging Data

To determine if a subjective valuation of effort better accounted for neural activity in vmPFC than an objective representation, we performed a Bayesian model selection analysis (Rosa et al., 2010). We began by creating an additional GLM that was identical to our original model *DV_sure_*(*G*,*S*), except in this model the parametric modulator corresponded to the difference in expected objective value of the effort options presented 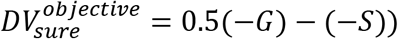. This GLM captured the null choice model (objective valuation of effort; *ρ* = 1).

We used the first level Bayesian estimation procedure in SPM12 to compute voxel-wise whole-brain log-model evidence maps for every participant and each model. To model inference at the group level we applied a random effects approach at every voxel of the log evidence data across the whole brain. We used this data to create exceedance probability maps (EPM) that allowed us to test which representation of effort cost, subjective or objective, was more likely to describe activity in vmPFC. The EPMs shown illustrate clusters of voxels at which subjective effort valuation has a greater Bayesian probability (*P* > 0.95) of describing the observed BOLD signal in vmPFC.

Additionally, in a similar manner to the analysis described above, we performed a second Bayesian model comparison, this time investigating if difference in value between the effort options or choice difficulty was more likely to describe activity in ACC. From the whole-brain log-evidence maps of each subject, comparing these two models, we were able to generate an EPM within the ACC ROI previously described.

## RESULTS

### Behavioral Representations of Subjective Effort Valuation

Comparison between reported and exerted effort levels during the recall phase showed a high degree of agreement, indicating that participants accurately perceived the objective effort levels (Fig. 2A shows the group recall results).

**Figure 2.**
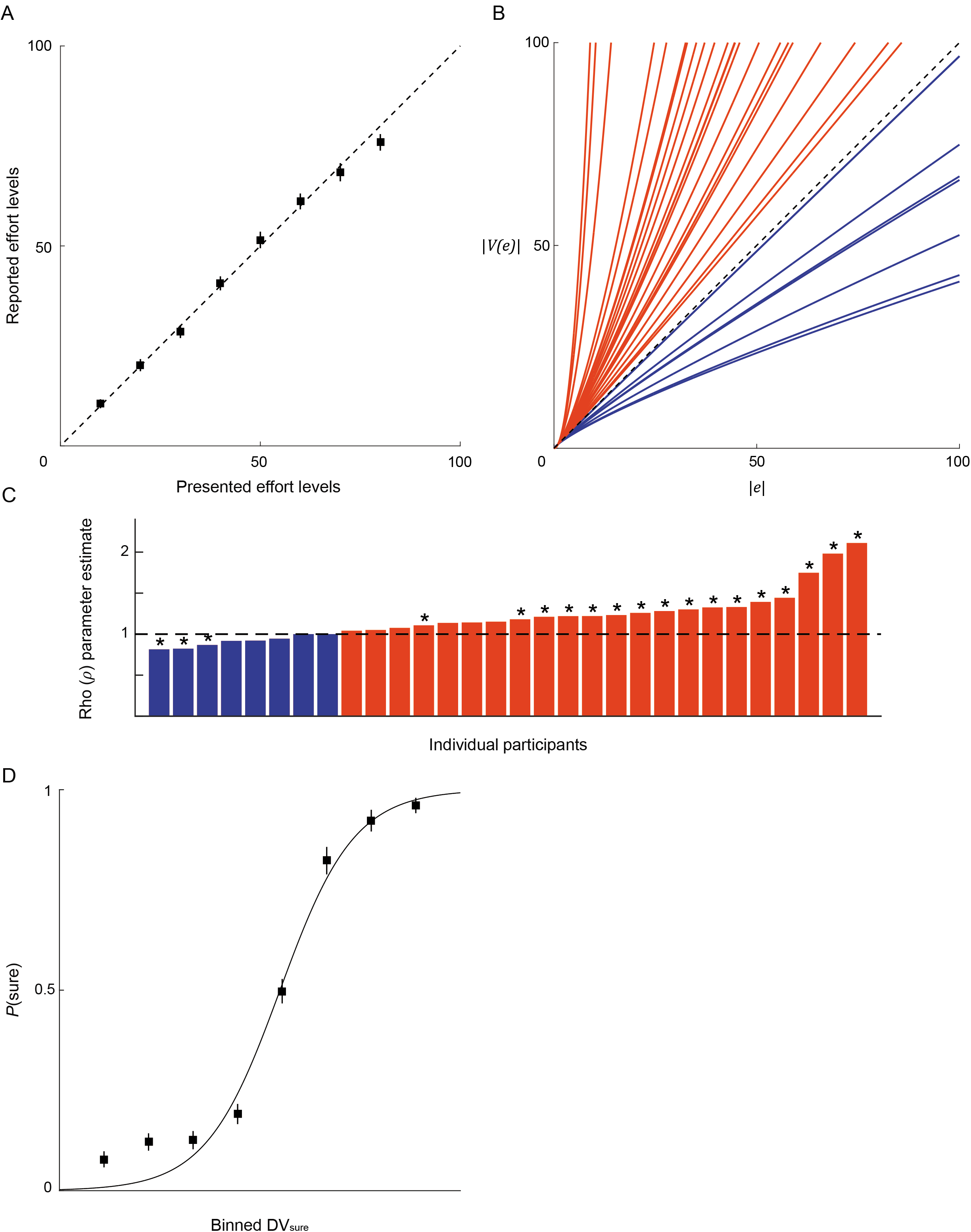
Behavioral representations of subjective effort cost. A. Results from the Recall phase of the experiment, showing the mean and standard error across all participants for the effort levels reported plotted against the those tested. The dashed line is included to indicate perfect recall of exerted effort.
B. The function used to model the subjective cost of effort in a choice. This function has the form *V*(*e*) = −(−*e*)^*ρ*^. Each curve represents an individual’s cost function for effort. The dashed line is included to indicate an objective valuation of effort (*ρ* = 1), with curves above this line representing that an incremental change in the effort level results in a greater subjective cost of that effort for higher effort levels.
C. Estimated *ρ* parameters at the participant level. Asterisks indicate a significant difference (p < 0.05) from the null hypothesis of objective valuation (*ρ* = 1) using a likelihood ratio test statistic.
D. Propensity to accept the sure option as a function of DV_sure_. DV_sure_ was partitioned into eight bins and the mean and standard error of the acceptance rate within each bin is displayed.

We characterized the subjectivity of participant *i’s* effort choices using a subjective cost function *V_i_*(*e*) = −(−*e*)^*ρ_i_*^, where *e* ≤ 0 and *V* is the subjective cost of an objective effort level *e*. *ρ_i_* is a participant-specific parameter that characterizes how an individual subjectively represents the effort level *e*. In this formulation, *ρ* is flexible enough to capture increasing, decreasing, or constant marginal changes in subjective effort valuation as absolute effort levels increase (Fig. 2B). The case where *ρ* = 1 indicates that a participant’s subjective effort cost coincides with absolute effort levels; *ρ* < 1 indicates decreasing sensitivity to changes in subjective effort cost as effort level increases; *ρ* > 1 indicates increasing sensitivity to changes in subjective effort cost as effort level increases.

Using the behavioral data, we performed a maximum likelihood estimation procedure to characterize each participant’s subjectivity of effort cost *ρ* and underlying consistency of choice *τ*. We found that participants exhibited mean parameter estimates of *ρ* = 1.20 (S.D. 0.30), *τ* = 0.23 (S.D. 0.26). A parameter recovery procedure found a significant correlation between parameters initially estimated and those recovered, suggesting that the participants’ decisions over the effort choice options yielded a precise estimation of *ρ* (see Supplementary Material for details). A likelihood ratio test statistic indicated that the majority of participants (*n* = 19) made choices that were inconsistent with a linear subjective effort function (*ρ* = 1), and the group exhibited subjectivity parameters that were significantly greater than 1 (*t*_29_ = 3.61, *p* < 0.001). We also computed a group level AIC using log-likelihood measures obtained from the MLE procedure and found AIC_objective_ = 4958 and AIC_subjective_ = 4552, further indicating that subjective valuation of effort best describes participants’ choices. Together, these results reveal that participants do not make effort decisions purely based on an objective valuation of effort, and that the majority of participants instead exhibited subjectivity of effort such that larger effort levels yielded increased marginal effort costs (Fig. 2C). These behavioral findings are consistent with previous studies that modeled subjectivity of effort valuation when trading effort for reward (Klein-Flügge et al. 2015; Klein-Flügge et al. 2016; Chong et al. 2017).

We performed a series of analyses to determine if the subjective effort parameters were related to other potential factors that could influence effort valuation. We found that participants’ effort subjectivity parameters were not significantly correlated with MVC (*r* = −0.08, *p* = 0.69), suggesting that subjective preferences for effort were not simply the byproduct of maximum strength. Additionally, effort subjectivity parameters did not correlate with measures of monetary subjective value (risk aversion: *r* = −0.21, *p* = 0.38; loss aversion: *r* = −0.17, *p* = 0.50), suggesting that individuals’ subjective preferences for reward are not related to their effort preferences and not simply a reflection of similar risk attitudes across decisions for different types of goods. Another possibility is that effort subjectivity parameters could be a reflection of the probability of success during the association phase – participants that were less successful at achieving the targeted exertions might find effort to be more costly and have higher *ρ* parameters. To test this possibility we examined the relationship between *ρ* and success rate during the association phase. Again, we did not find a significant relationship between the two (*r* = −3.54 × 10^−4^, *p* = 0.99), suggesting that *ρ* is not driven by the probability of success during association.

Next we investigated the relationship between subjective effort valuation and decision difficulty. We found that participants’ decisions revealed that the difference in subjective utility between the two effort options *DV_sure_*(*G*,*S*) = 0.5(−*G*)^*ρ*^ − (− *S*)^*ρ*^ sampled the range of option rejection/acceptance (Figure 2D). With this in mind, choice difficulty should be greatest when the effort options are most similar (when -|DV_sure_| is near 0), and least difficult when the options are most dissimilar. Accordingly, response time will be longest when -|DV_sure_| is small and shortest when choices are easiest. Consistent with this idea, we found that model-free (log(response time)) and model-based (-|DV_sure_|) measures of choice difficulty were significantly positively correlated with one another (average Pearson’s correlation *r* = 0.05; one tailed: *t*_29_ = 1.80, *p* = 0.04). Critically, these measures of choice difficulty were not correlated with the difference in subjective utility between the two options (DV_sure_ and log(reaction time): average Pearson’s correlation *r* = −0.41 × 10^−2^, *t*_29_ = −0.17, *p* = 0.87; DV_sure_ and -|DV_sure_|: average Pearson’s correlation *r* = −0.03, *t*_29_ = −0.25, *p* = 0.80). This orthogonalization between the subjective utility of effort and choice difficulty allowed us to identify the neural signals associated with each computational variable.

### vmPFC Encodes Subjective Valuation of Effort

To test our neural hypothesis that subjective effort valuation is encoded in vmPFC, we estimated a general linear model (GLM) in SPM12 of the blood-oxygenation level dependent (BOLD) activity of the whole-brain during the choice phase. This model included parametric modulators at the time of choice, corresponding to both the difference in value between the sure and gamble options DV_sure_ and choice difficulty as indexed by log(reaction time). DV_sure_ was defined by transforming the effort options under consideration using the effort subjectivity parameter *ρ* estimated from each individual participant’s behavior (see Materials and Methods for details). This formulation allowed us to isolate brain regions that encoded subjective valuation of effort and choice difficulty at the time of decision.

We found that BOLD signal in vmPFC was significantly positively correlated with DV_sure_ (ROI analysis of vmPFC: *t*_29_ = 2.27, *p* = 0.03; Fig. 3A, B). As DV_sure_ increased, vmPFC activity significantly increased suggesting that this region encoded the difference in subjective effort value of the two options. The areas of vmPFC identified largely overlapped those found in studies of appetitive and aversive valuation (Bartra et al. 2013; Clithero and Rangel 2014; O’Doherty 2014). Notably we did not find a significant effect of choice difficulty in vmPFC (ROI analysis of vmPFC: *t*_29_ = −1.47, *p* = 0.15).

**Figure 3.**
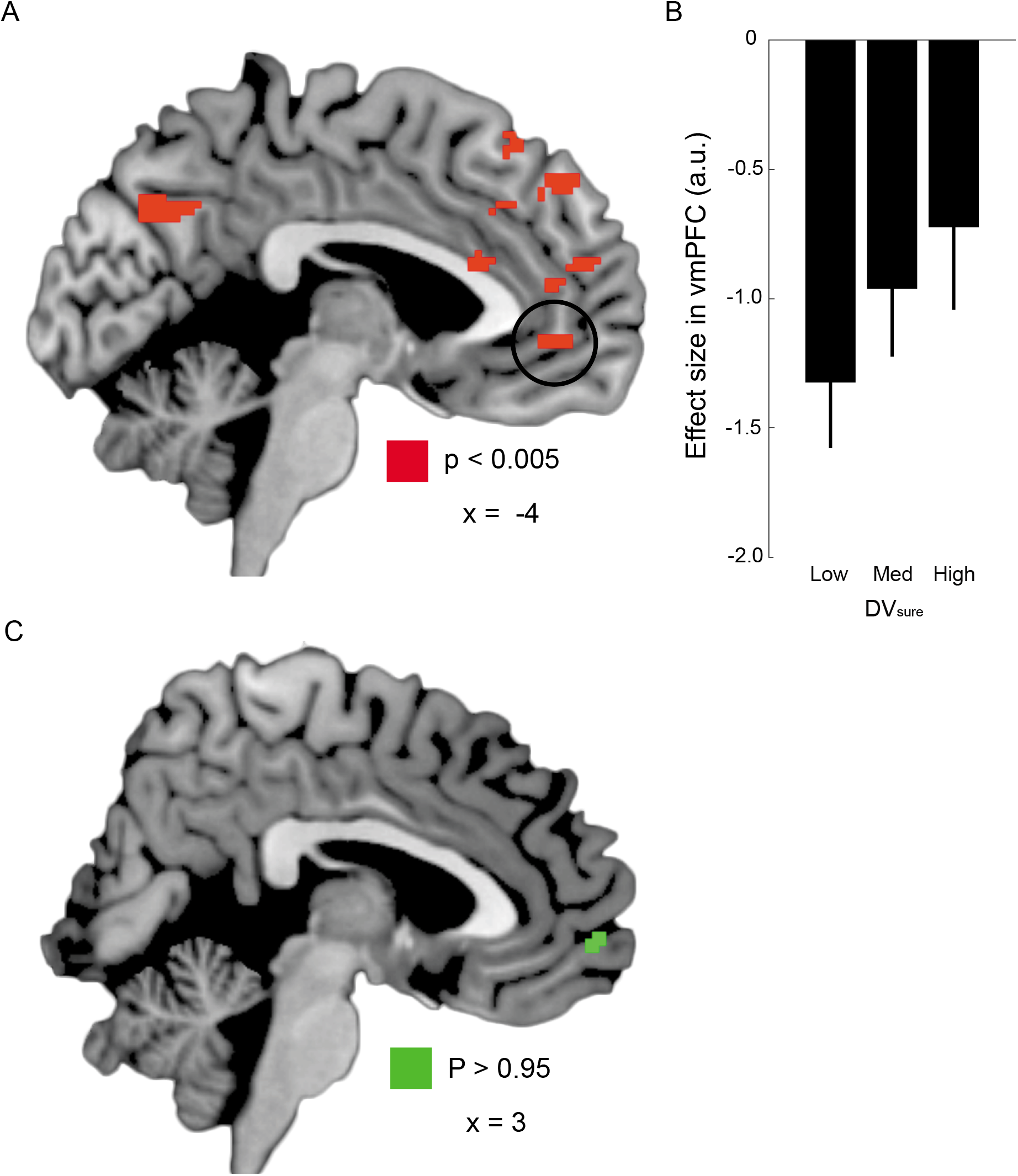
vmPFC encodes subjective effort valuation. A. A region of vmPFC in which BOLD activity was positively correlated with the decision value of the sure option at the time of choice, with peak activity at Montreal Neurological Institute (MNI) coordinates (x, y, z) = [−4, 46, −2]. The contrast shown in red was obtained at p < 0.005 (uncorrected) with a 10 voxel extent threshold. This contrast is significant at p < 0.05, small volume corrected in an independent vmPFC ROI.
B. BOLD effect size within a 5 mm sphere centered at peak activity in vmPFC was positively correlated with the difference in utility between the two options (DV_sure_). This plot is not used for statistical inference (which was performed using an independent ROI analysis); it is show shown solely to illustrate the trend of the BOLD signal in vmPFC.
C. Exceedance probability map (EPM) resulting from the Bayesian model comparison of objective and subjective effort valuation models. Voxels shown in green (*n* = 16) indicate locations in the brain where the probability that subjective effort describes the BOLD activity is greater than 0.95, supporting the finding that subjective effort cost best describes activity in vmPFC.

In a separate test, to confirm that activity in vmPFC during effort choices was best described by representing options subjectively as opposed to objectively (*ρ* = 1), we generated exceedance probability maps (EPMs) for the imaging model described above as well as a null model representing objective effort valuation 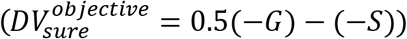 (Rosa et al. 2010). Using these probabilistic brain maps we were able to evaluate the likelihood that areas of vmPFC better represented subjective effort costs as opposed to objective effort costs. We found a cluster of voxels in vmPFC, (16 voxels, *P* > 0.95), illustrating that activity in this region is best described by a subjective rather than objective model of effort costs (Fig. 3C).

### ACC Encodes Choice Difficulty

To test our hypothesis that choice difficulty is encoded in ACC we used the previously described imaging model to identify brain activity that was correlated with increasing choice difficulty as indexed by log(response time). We found that choice difficulty was significantly positively correlated with activity in ACC (ROI analysis of ACC: *t*_29_ = 2.07, *p* = 0.04; Figure 4A, B), consistent with previous studies that have shown that this region encodes choice difficulty during neuroeconomic choice (Shenhav et al. 2014; Shenhav et al. 2016).

**Figure 4.**
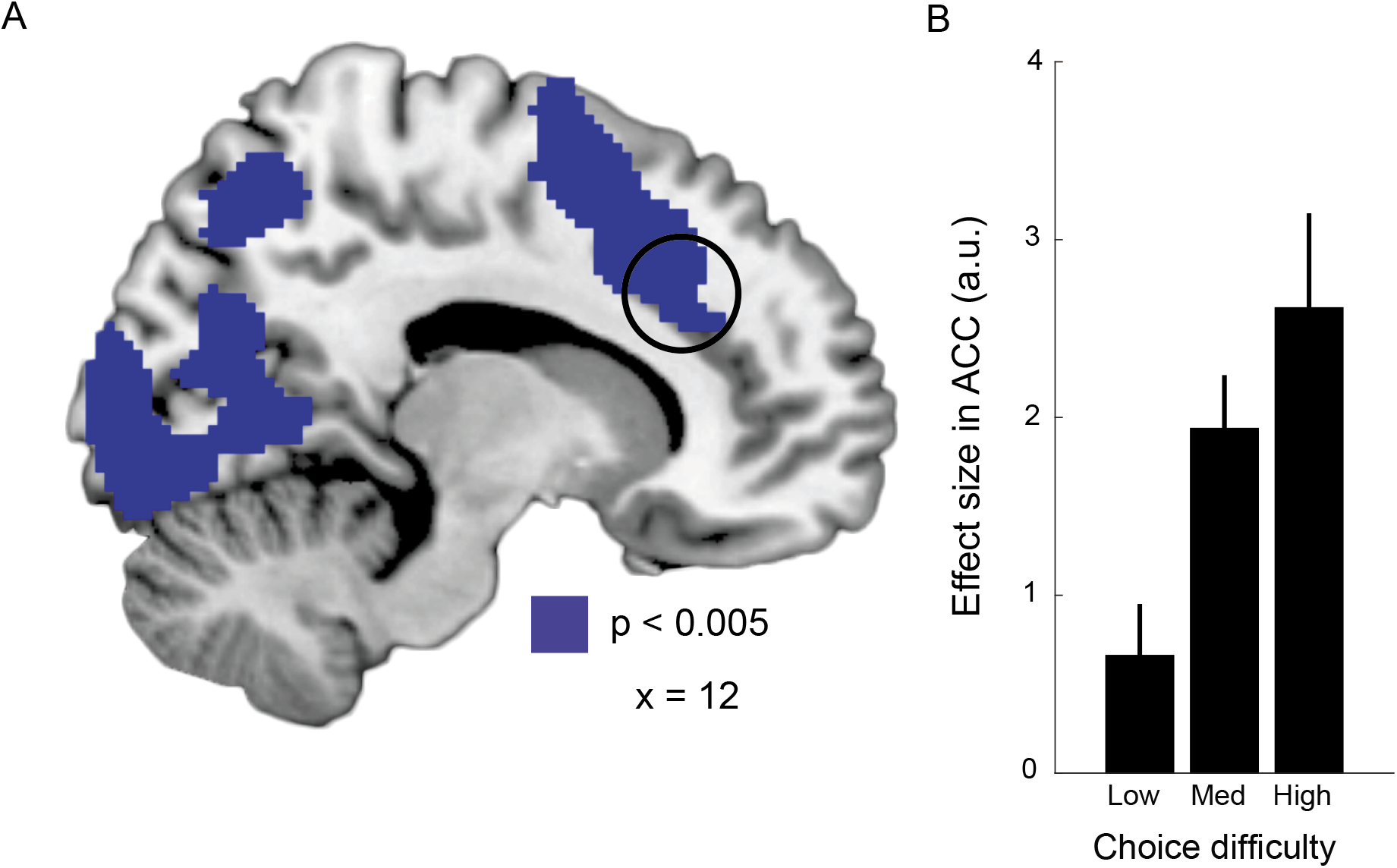
ACC encodes choice difficulty. A. Regions of the brain in which BOLD activity was positively correlated with choice difficulty at the time of choice with peak activity at Montreal Neurological Institute (MNI) coordinates (x, y, z) = [−2, 14, 42]. The contrast shown in blue was obtained at p < 0.005 (uncorrected) with a 10 voxel extent threshold. This contrast is significant at p < 0.05, small volume corrected in an independent ACC ROI.
B. BOLD effect size within a 5 mm sphere centered at peak activity in was positively correlated choice difficulty. This plot is not used for statistical inference (which was performed using an independent ROI analysis); it is show shown solely to illustrate the trend of the BOLD signal in ACC.

We also searched for effort value signals in ACC, as have been reported in previous studies, however even decreasing our contrast significance level to p < 0.01 did not reveal any significant activation in our ACC effort ROI. To further test if ACC encoded representations of subjective effort valuation, we performed a formal ROI analysis of ACC. This analysis did not find activation in ACC related to DV_sure_ that reached significance (ROI analysis of ACC: *t*_29_ = 1.66 *p* = 0.11). To directly confirm that activity in ACC was best described by choice difficulty as opposed to DV_sure_ we generated an additional EPM incorporating both choice difficulty and difference in effort value. This analysis revealed that activity in ACC was better described by choice difficulty rather than DV_sure_ (ROI analysis of ACC: average probability across all voxels *P* = 0.93, SD = 0.10).

It should be noted that in the aforementioned analyses we used a noisy model-free representation of choice difficulty: log(response time). We also performed a second analysis, similar to the first, except we included a model-based measure of choice difficulty (-|DV_sure_|) along with log(response time) and DV_sure_. Again, we found significant activations in ACC positively correlated with -|DV_sure_| (ROI analysis of ACC: one-tailed: *t*_29_ = 2.04, *p* = 0.03). These results indicate that even when accounting for behavioral noise captured by response time, choice difficulty elicits significant activations in ACC.

## DISCUSSION

We used a novel effort choice paradigm in which participants were presented with prospective effort options under uncertainty, which were independent of reward and orthogonal to choice difficulty. This paradigm allowed us to isolate neural signals related to subjective valuation of effort that were not contingent on reward or concomitant with choice difficulty. Behaviorally, we found that the average individual’s subjective effort cost exhibited increasing marginal costs as effort increased. Neurally, we found that activity in vmPFC was related to the subjective valuation of prospective effort, while ACC activity was related to choice difficulty. These results suggest that vmPFC encodes the subjective costs that underlie choices involving physical effort, and ACC activity is related to the cognitive control required at the time of choice.

Previous studies of effort cost have focused on the trade-offs between prospective effort and reward, similar to the natural choices we make in everyday life (Croxson et al. 2009; Prévost et al. 2010; Kurniawan et al. 2010; Kurniawan et al. 2013; Skvortsova et al. 2014; Klein-Flügge et al. 2016; Chong et al. 2017), and have suggested that in these contexts ACC encodes effort cost. However these studies were not designed to separate effort costs from reward and instead focused on the integration of both of these utilities to compute a decision. In our paradigm, however, we took a reductionist scientific approach, isolating effort from reward in order to provide a computational description of how subjective valuation of effort is encoded in the brain. While our choice paradigm is not as naturalistic as the effort/reward trade-offs made in daily life, such an approach is valuable because it affords us a deeper understanding of an individual component (in this case effort valuation) that shapes more complex decisions. Our paradigm revealed that activity in vmPFC was better represented by subjective, rather than objective, valuation of prospective effort. This is consistent with previous research in an intertemporal choice setting that finds this region better encodes subjective monetary values compared to objective monetary values (Kable and Glimcher 2007). Moreover our findings are consistent with the idea that vmPFC encodes a general subjective value signal that subserves effort decisions, similar to the subjective value signals that have been previously reported for a variety of appetitive and aversive stimuli (Bartra et al. 2013; Clithero and Rangel 2014; O’Doherty 2014).

Notably, we did not find activity in ACC in our fMRI analysis of subjective effort valuation, but instead found that it was related to choice difficulty, consistent with previous studies of neuroeconomic choice that experimentally separated prospective value and choice difficulty. There is an ongoing debate regarding the role of ACC in decision-making and whether it encodes decision values or variables related to cognitive control (e.g., choice difficulty) (Shenhav et al. 2016; Kolling et al. 2016; Ebitz and Hayden 2016). With this debate in mind it has been proposed that when studying valuation it is important to design studies that are capable of separating the two. Aside from one previous study of effort-based decision-making, none have controlled for the relationship between these two variables. The one study that attempted to account for choice difficulty reported that it did not describe activity in ACC, however that design did not span the full space of prospective effort to elicit a full range of choice preferences (Chong et al. 2017) making it challenging to truly dissociate signals related to valuation and choice difficulty. Our paradigm was designed to span the full space of effort valuation and choice preference, and in doing so we found that ACC activity was associated with choice difficulty, as opposed to effort valuation. Our result of effort based choice difficulty signals in ACC, taken together with previous studies of foraging (Shenhav et al. 2014; Shenhav et al. 2016) are consistent with ACC’s role in cognitive control during decision-making. In particular, our findings align with conflict monitoring theories that suggest ACC tracks the level of indifference in decision-making tasks because higher indifference requires increased cognitive control (Botvinick et al. 2001; Botvinick 2007). It is important to mention that it is also possible that such conflict signals could be a byproduct of ACC comparing the values of the options presented (Hare et al. 2011), a property that would be consistent with conflict monitoring theories.

It has also been proposed that the ACC activity found in previous studies could be indicative of a multiplexing node that combines action/reward values (Hayden and Platt 2010; Shenhav et al. 2013; Klein-Flügge et al. 2016) and serves as a gateway that informs the motor system to act for reward (Cai and Padoa-Schioppa 2012). Thus we cannot rule out the possibility that our lack of observed subjective effort value signals in ACC could also be due to the fact that our study was designed to isolate neural representations of subjective effort valuation that were *independent* of (and not multiplexed with) reward.

While it is important to interpret null results in fMRI imaging with caution, our results suggest that in the context of effort-based decision-making, in which rewards are not present and choice difficulty is controlled, the ACC better represents choice difficulty than effort valuation. While this finding is in contrast to previous studies of effort/reward tradeoffs that have implicated ACC in this process, it is important to stress that our study attempted to isolate effort valuation irrespective of reward and choice difficulty, and thus was quite different from those previous investigations. Future studies will be needed to determine, if when controlling for choice difficulty in more naturalistic paradigms involving effort reward trade-offs, ACC activity is still best described by difficulty and vmPFC by effort valuation. Notably, in the ongoing debate about ACC function during economic choice, it has been suggested that sub-regions within ACC might simultaneously encode signals about valuation and choice difficulty (Kolling et al., 2016a; Kolling et al., 2016b), and in this vein it is possible that ACC could encode both effort costs and choice difficulty (although in this study we did not find evidence in support of this idea).

In this study we focused on characterizing the subjective valuation of physical effort in the form of grip force. However, an individual’s subjective effort costs could vary across types of effort (i.e., walking, arm movements, or even cognitive effort) in a similar fashion to how individuals exhibit different subjective values for different types of goods (Chib et al. 2009; Levy and Glimcher 2011). Moreover, just as the subjective value of rewards can be modulated by the state of an individual, subjective costs of effort could also be influenced by state. For example, individuals having undergone physical or cognitive fatigue or training might exhibit modified representations of subjective effort cost. Furthermore, it is possible that the subjectivity of different types of effort may exhibit similar trait-like consistency over time, as has been reported in studies of subjective valuation of money (Ohmura et al. 2006; Kable and Glimcher 2007; Ballard and Knutson 2009).

Characterization of subjective effort costs will provide an understanding of why some people find certain tasks to be very effortful while others complete them with ease. Such knowledge could be used to design incentive mechanisms that account for perceptions of effortfulness to maximize employees’ performance. Insights into these preferences may aid in the development of more efficacious individual-specific behavioral mechanisms that enhance motivational output and effort exertion in a variety of everyday tasks.

## ACKNOWLEDGEMENTS

The authors declare no competing financial interests.

Acknowledgements: We thank D. McNamee and V. Stuphorn for their insightful comments on this study, and C. Frydman for his help with the design and analysis. This work was supported by the Eunice Kennedy Shriver National Institute Of Child Health & Human Development of the National Institutes of Health under Award Number K12HD073945 to V.S.C. J.G. was supported by the National Defense Science and Engineering Graduate Fellowship. The authors declare no competing financial interests.

## REFERENCES

Abeler J, Falk A, Goette L, Huffman D (2011) Reference Points and Effort Provision. Am Econ Rev 101:470–492.

Augenblick N, Niederle M, Sprenger C (2015) Working Over Time: Dynamic Inconsistency in Real Effort Tasks*. Q J Econ:qjv020.

Ballard K, Knutson B (2009) Dissociable neural representations of future reward magnitude and delay during temporal discounting. NeuroImage 45:143–150.

Bartra O, McGuire JT, Kable JW (2013) The valuation system: A coordinate-based meta-analysis of BOLD fMRI experiments examining neural correlates of subjective value. NeuroImage 76:412–427.

Botvinick MM, Braver TS, Barch DM, Carter CS, Cohen JD (2001) Conflict monitoring and cognitive control. Psychol Rev 108(3):624–652.

Botvinick MM (2007) Conflict monitoring and decision making: Reconciling two perspectives on anterior cingulate function. Cogn Affect Behav Neurosci 7:356–366.

Brainard DH (1997) The Psychophysics Toolbox. Spat Vis 10:433–436.

Cai X, Padoa-Schioppa C (2012) Neuronal Encoding of Subjective Value in Dorsal and Ventral Anterior Cingulate Cortex. J Neurosci 32:3791–3808.

Camerer C, Loewenstein G, Prelec D (2005) Neuroeconomics: How Neuroscience Can Inform Economics. J Econ Lit 43:9–64.

Chib VS, De Martino B, Shimojo S, O’Doherty JP (2012) Neural Mechanisms Underlying Paradoxical Performance for Monetary Incentives Are Driven by Loss Aversion. Neuron 74:582–594.

Chib VS, Rangel A, Shimojo S, O’Doherty JP (2009) Evidence for a Common Representation of Decision Values for Dissimilar Goods in Human Ventromedial Prefrontal Cortex. J Neurosci 29:12315–12320.

Chib VS, Shimojo S, O’Doherty JP (2014) The Effects of Incentive Framing on Performance Decrements for Large Monetary Outcomes: Behavioral and Neural Mechanisms. J Neurosci 34:14833–14844.

Chong TT-J, Apps M, Giehl K, Sillence A, Grima LL, Husain M (2017) Neurocomputational mechanisms underlying subjective valuation of effort costs. PLOS Biol 15:e1002598.

Clithero JA, Rangel A (2014) Informatic parcellation of the network involved in the computation of subjective value. Soc Cogn Affect Neurosci 9:1289–1302.

Croxson PL, Walton ME, O’Reilly JX, Behrens TEJ, Rushworth MFS (2009) Effort-Based Cost–Benefit Valuation and the Human Brain. J Neurosci 29:4531–4541.

Deichmann R, Gottfried JA, Hutton C, Turner R (2003) Optimized EPI for fMRI studies of the orbitofrontal cortex. Neuroimage 19:430–441.

DellaVigna S, Paserman MD (2005) Job Search and Impatience. J Labor Econ 23:527–588.

Dishman RK (1991) Increasing and maintaining exercise and physical activity. Behav Ther 22:345–378.

Ebitz RB, Hayden BY (2016) Dorsal anterior cingulate: a Rorschach test for cognitive neuroscience. Nat Neurosci 19:1278–1279.

Floresco SB, Onge JRS, Ghods-Sharifi S, Winstanley CA (2008) Cortico-limbic-striatal circuits subserving different forms of cost-benefit decision making. Cogn Affect Behav Neurosci 8:375–389.

Frydman C, Camerer C, Bossaerts P, Rangel A (2011) MAOA-L carriers are better at making optimal financial decisions under risk. Proc R Soc Lond B Biol Sci 278:2053–2059.

Hare TA, Schultz W, Camerer CF, O’Doherty JP, Rangel A (2011) Transformation of stimulus value signals into motor commands during simple choice. Proc Natl Acad Sci 108:18120–18125.

Hayden BY, Heilbronner SR (2014) All that glitters is not reward signal. Nat Neurosci 17:1142–1144.

Hayden BY, Platt ML (2010) Neurons in Anterior Cingulate Cortex Multiplex Information about Reward and Action. J Neurosci 30:3339–3346.

Hillman KL, Bilkey DK (2012) Neural encoding of competitive effort in the anterior cingulate cortex. Nat Neurosci 15:1290–1297.

Holt CA, Laury SK (2002) Risk Aversion and Incentive Effects. Am Econ Rev 92:1644–1655.

Kable JW, Glimcher PW (2007) The neural correlates of subjective value during intertemporal choice. Nat Neurosci 10:1625–1633.

Kahneman D, Tversky A (1979) Prospect Theory: An Analysis of Decision under Risk. Econometrica 47:263–291.

Klein-Flügge MC, Kennerley SW, Friston K, Bestmann S (2016) Neural Signatures of Value Comparison in Human Cingulate Cortex during Decisions Requiring an Effort-Reward Trade-off. J Neurosci 36:10002–10015.

Klein-Flügge MC, Kennerley SW, Saraiva AC, Penny WD, Bestmann S (2015) Behavioral Modeling of Human Choices Reveals Dissociable Effects of Physical Effort and Temporal Delay on Reward Devaluation. PLOS Comput Biol 11:e1004116.

Kolling N, Behrens T, Wittmann M, Rushworth M (2016a) Multiple signals in anterior cingulate cortex. Curr Opin Neurobiol 37:36–43.

Kolling N, Wittmann MK, Behrens TEJ, Boorman ED, Mars RB, Rushworth MFS (2016b) Value, search, persistence and model updating in anterior cingulate cortex. Nat Neurosci 19:1280–1285.

Kriegeskorte N, Simmons WK, Bellgowan PSF, Baker CI (2009) Circular analysis in systems neuroscience: the dangers of double dipping. Nat Neurosci 12:535–540.

Kurniawan IT, Guitart-Masip M, Dayan P, Dolan RJ (2013) Effort and Valuation in the Brain: The Effects of Anticipation and Execution. J Neurosci 33:6160–6169.

Kurniawan IT, Seymour B, Talmi D, Yoshida W, Chater N, Dolan RJ (2010) Choosing to Make an Effort: The Role of Striatum in Signaling Physical Effort of a Chosen Action. J Neurophysiol 104:313–321.

Levy DJ, Glimcher PW (2011) Comparing Apples and Oranges: Using Reward-Specific and Reward-General Subjective Value Representation in the Brain. J Neurosci 31:14693–14707.

O’Doherty JP (2014) The problem with value. Neurosci Biobehav Rev 43:259–268.

Ohmura Y, Takahashi T, Kitamura N, Wehr P (2006) Three-month stability of delay and probability discounting measures. Exp Clin Psychopharmacol 14:318–328.

Prévost C, Pessiglione M, Météreau E, Cléry-Melin M-L, Dreher J-C (2010) Separate Valuation Subsystems for Delay and Effort Decision Costs. J Neurosci 30:14080–14090.

Rangel A, Camerer C, Montague PR (2008) A framework for studying the neurobiology of value-based decision making., Neuroeconomics: The neurobiology of value-based decision-making. Nat Rev Neurosci Nat Rev Neurosci 9, 9:545, 545–556.

Rosa MJ, Bestmann S, Harrison L, Penny W (2010) Bayesian model selection maps for group studies. NeuroImage 49:217–224.

Rudebeck PH, Behrens TE, Kennerley SW, Baxter MG, Buckley MJ, Walton ME, Rushworth MFS (2008) Frontal Cortex Subregions Play Distinct Roles in Choices between Actions and Stimuli. J Neurosci 28:13775–13785.

Shenhav A, Botvinick MM, Cohen JD (2013) The expected value of control: an integrative theory of anterior cingulate cortex function. Neuron 79:217–240.

Shenhav A, Straccia MA, Botvinick MM, Cohen JD (2016) Dorsal anterior cingulate and ventromedial prefrontal cortex have inverse roles in both foraging and economic choice. Cogn Affect Behav Neurosci 16:1127–1139.

Shenhav A, Straccia MA, Cohen JD, Botvinick MM (2014) Anterior cingulate engagement in a foraging context reflects choice difficulty, not foraging value. Nat Neurosci 17:1249–1254.

Skvortsova V, Palminteri S, Pessiglione M (2014) Learning To Minimize Efforts versus Maximizing Rewards: Computational Principles and Neural Correlates. J Neurosci 34:15621–15630.

Sniehotta FF, Scholz U, Schwarzer R (2005) Bridging the intention–behaviour gap: Planning, self-efficacy, and action control in the adoption and maintenance of physical exercise. Psychol Health 20:143–160.

Sokol-Hessner P, Camerer CF, Phelps EA (2012) Emotion regulation reduces loss aversion and decreases amygdala responses to losses. Soc Cogn Affect Neurosci:nss002.

Sokol-Hessner P, Hsu M, Curley NG, Delgado MR, Camerer CF, Phelps EA (2009) Thinking like a trader selectively reduces individuals’ loss aversion. Proc Natl Acad Sci 106:5035–5040.

Walton ME, Bannerman DM, Rushworth MFS (2002) The Role of Rat Medial Frontal Cortex in Effort-Based Decision Making. J Neurosci 22:10996–11003.

Walton ME, Groves J, Jennings KA, Croxson PL, Sharp T, Rushworth MFS, Bannerman DM (2009) Comparing the role of the anterior cingulate cortex and 6-hydroxydopamine nucleus accumbens lesions on operant effort-based decision making. Eur J Neurosci 29:1678–1691.

Westbrook A, Braver TS (2015) Cognitive effort: A neuroeconomic approach. Cogn Affect Behav Neurosci 15:395–415.

